# In silico MS/MS prediction for peptidoglycan profiling uncovers novel anti-inflammatory peptidoglycan fragments of the gut microbiota

**DOI:** 10.1101/2023.10.08.561446

**Authors:** Jeric Mun Chung Kwan, Yaquan Liang, Evan Wei Long Ng, Ekaterina Sviriaeva, Chenyu Li, Yilin Zhao, Xiao-Lin Zhang, Xue-Wei Liu, Sunny H. Wong, Yuan Qiao

## Abstract

Peptidoglycan is an essential exoskeletal polymer present across all bacteria. The gut microbiota-derived peptidoglycan fragments (PGNs) are increasingly recognized as key effector molecules that impact host biology, offering attractive yet untapped potential to combat microbiome-associated diseases in humans. Unfortunately, comprehensive peptidoglycan profiling of gut bacteria has been hampered by the lack of a robust and automated analysis workflow. Currently, PGN identification still relies on manual deconvolutions of acquired tandem mass spectrometry (MS/MS) data, which are highly laborious and inconsistent. Recognizing the unique sugar and amino acid makeup of bacterial peptidoglycan and guided by the experimental MS/MS fragmentation patterns of known PGNs, we developed a computational tool PGN_MS2 that reliably simulates MS/MS spectra of PGNs. Integrating PGN_MS2 into the customizable *in silico* PGN database, we built an open-access PGN MS library of predicted MS/MS spectra for all molecules in the user-defined *in silico* PGN search space. With this library, automated searching and spectral matching can be used to identify PGN. We then performed comprehensive peptidoglycan profiling for several gut bacteria species, revealing distinct PGN structural features that may be implicated in microbiota-host crosstalk. Strikingly, the probiotic *Bifidobacterium* spp. has an exceedingly high proportion of anhydro-PGNs, which exhibit anti-inflammatory effects *in vitro*. We further identified MltG and RfpB homologs in *Bifidobacterium* as lytic transglycosylases (LTs), which demonstrate distinct substrate preferences to produce anhydro-PGNs. Overall, our novel PGN_MS2 prediction tool contributes to the robust and automated peptidoglycan analysis workflow, advancing efforts to elucidate the structures and functions of gut microbiota-derived PGNs in the host.

## Introduction

All bacteria possess a peptidoglycan layer. As an essential exoskeletal polymer that surrounds the bacterial cytoplasmic membrane, peptidoglycan protects bacterial cells against internal turgor pressure and also serves as a scaffold for other cell surface proteins and polymers.(1) Disruption of the peptidoglycan layer’s integrity can effectively lead to bacterial lysis and death. Peptidoglycan modifications in certain bacteria are known to confer resistance against peptidoglycan-targeting antibiotics as well as lytic enzymes, highlighting the biological significance of peptidoglycan structural features for bacterial survival.(2–6)

Apart from a structural role, bacterial peptidoglycan also participates in diverse intra- and inter-kingdom signalling.(7, 8) Soluble peptidoglycan fragments, also known as PGNs or muropeptides, are continuously generated by bacteria during growth and released into the milieu, exerting a broad-range impact on different organisms.(9) In the context of the human gut microbiota, the trillions of resident bacteria produce a multitude of PGNs in the gut niche,(10) which can disseminate into host systemic circulation during steady-state conditions,(11) influencing host biology including autoimmunity, brain development, appetite, and body temperature, as well as efficacies of cancer immunotherapy.(12–15) Remarkably, subtle structural changes in PGNs can significantly alter their biological activities in hosts.(16) Thus, profiling peptidoglycan compositions and characteristics in gut bacteria is of paramount importance to facilitate studies of gut microbiota-derived PGNs in hosts.

While the chemical makeup of peptidoglycan is largely conserved, the exact compositions and structural modifications of peptidoglycan are highly variable across bacteria and under different environmental conditions (**Figure 1**).(1, 17, 18) In general, the ‘*glycan*’ component of peptidoglycan consists of alternating units of *N*-acetylglucosamine (GlcNAc, or herein NAG) and *N*-acetylmuramic acid (MurNAc, or herein NAM) linked via *β*-1,4-glycosidic bonds; the ‘*peptido*’ portion refers to the short stem pentapeptide connected onto the lactoyl group of each NAM, which has the common sequence of L-Ala_1_-γ-D-Glu/isoGln_2_-AA_3_-D-Ala_4_-D-Ala_5_, with AA_3_ being either L-Lys attached to a lateral bridge peptide (that is specific to each bacterial species) or a non-proteogenic diamino acid such as meso-diaminopimelic acid (mDAP) (**Figure 1A**). These stem peptides on adjacent glycan strands can further react to form isopeptide bonds via 3-4 or 3-3 crosslinks, thereby strengthening the peptidoglycan layer (**Figure 1C**). Furthermore, a great deal of the structural diversity in peptidoglycan comes from the cell wall remodelling process, where bacterial enzymes catalyze specific reactions at distinct positions in peptidoglycan to generate new structural moieties, such as modifications of the glycan backbone, trimming of pentapeptides to shorter stems, and incorporation of non-canonical D-amino acids (NCDAA) into the stem peptide (**Figure 1A-B**).(19) While most insights on peptidoglycan structural diversity were gained from analyses of the model bacterial organisms, our knowledge of the scope and variety of peptidoglycan in the gut microbiota is much in infancy. Recognizing the biological significance of peptidoglycan modifications, we seek to develop a robust and automated workflow for the characterization of peptidoglycan compositions and structural features in any bacteria of interest, such as gut bacteria.

**Figure 1.**
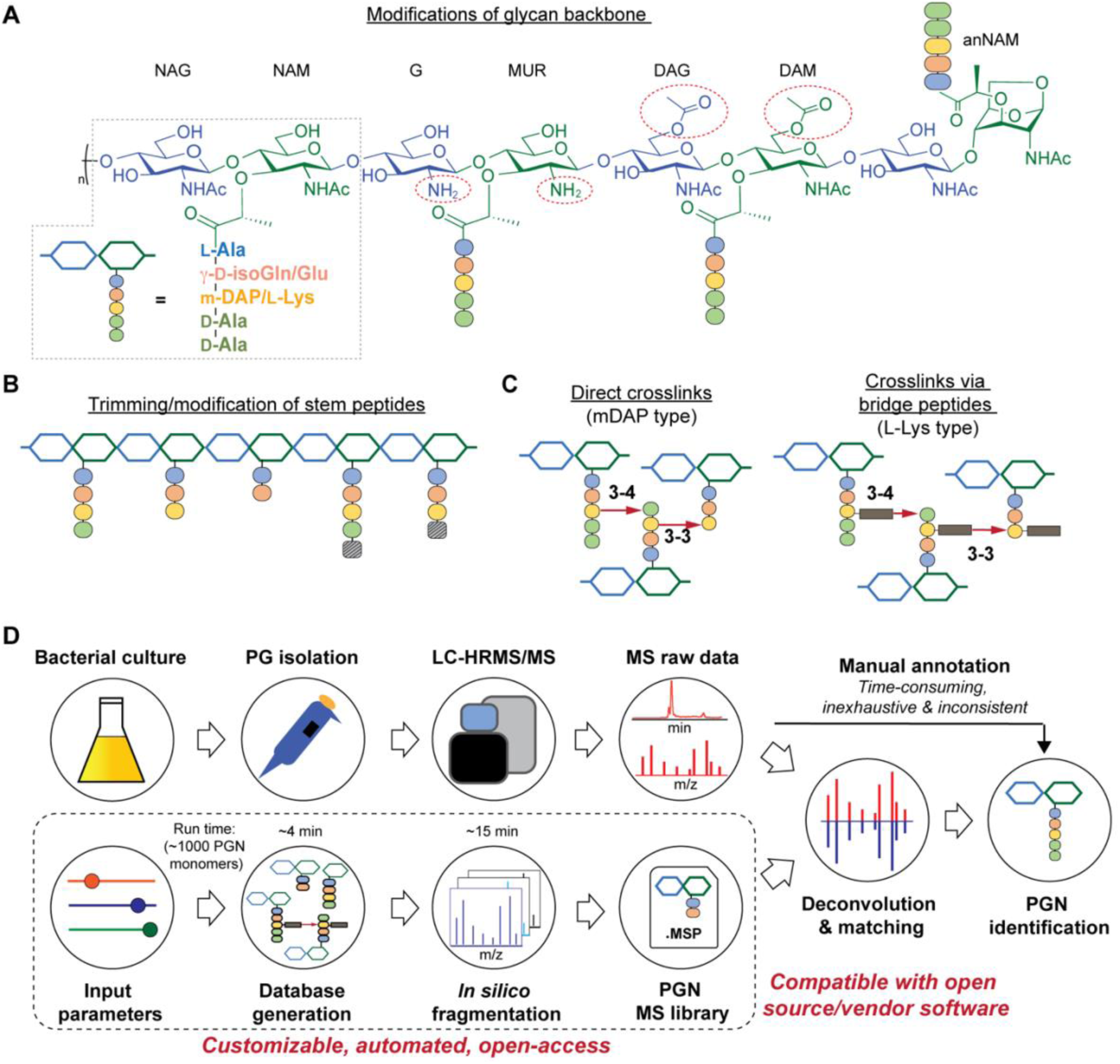
Schematic representations of bacterial peptidoglycan compositions (**A-C**) and our *in silico* peptidoglycan fragment (PGN) library advance the current analysis pipeline (**D**). (**A**) Peptidoglycan is composed of repeating units of disaccharide-muropeptides, which are *N-*acetylglucosamine (NAG)-*N*-acetylmuramic acid (NAM) with a stem peptide. The typical sequence of the stem peptide is L-Ala-γ-D-Glu-mDAP/L-Lys-D-Ala-D-Ala, where a species-specific bridge peptide is connected onto L-Lys. Moreover, the glycan backbone can exhibit various structural modifications. For instance, *N*-deacetylation on C2 for NAG and NAM yields glucosamine (G) and muramic acid (MUR), respectively; *O*-acetylation on C6 for NAG and NAM yields di-acetylated glucosamine (DAG) and di-acetylated muramic acid (DAM), respectively. 1,6-anhydro-*N-*acetylmuramic acid termini (anNAM) via intramolecular dehydration reaction catalysed by lytic transglycosylases (LTs). Glu: glutamate, isoGln: iso-glutamine; mDAP: meso-diaminopimelic acid. (**B**) Remodelling of peptidoglycan may result in trimming of the stem peptides and/or incorporation of non-canonical D-amino acids (represented by a striped box). (**C**) Adjacent stem peptides are crosslinked via direct crosslinks (left, for mDAP-containing peptidoglycans) or indirect, bridged crosslinks (right, for L-Lys-containing peptidoglycans). The 3-4 and 3-3 crosslinks refer to the positions of the crosslinked amino acids in the donor and acceptor stem peptides. (**D**) Current workflow of bacterial PGN analysis. Manual deconvolution of LC-MS/MS spectra for structural determination is the bottleneck. We present PGN_MS2, which creates a PGN database with user-provided parameters and automatically generates the *in silico* predicted MS/MS spectrum for each compound. The MS/MS spectra are compiled into a library (.msp) that is open-access and compatible with mass spectra analysis software for automated deconvolution of PGNs by *m/z*, isotopic pattern, and spectral similarity from the LC-MS/MS raw data.

There are significant gaps in the current workflow of bacterial peptidoglycan analysis, with the widely adopted experimental procedure developed >30 years ago.(20) Briefly, the peptidoglycan polymer (i.e., sacculi) isolated from bacteria is digested with a muramidase (e.g., lysozyme) that hydrolyzes the NAM-*β*-1,4-NAG linkages along the peptidoglycan backbone, generating soluble PGNs that are disaccharide-containing muropeptides in nature.(21) The collection of these soluble PGNs is then subjected to high-performance liquid chromatography-tandem mass spectrometry (HPLC-MS/MS) analysis for structural characterization and profiling (**Figure 1D**). Improvements in HPLC-MS/MS instrumentation such as higher resolution and faster scanning rate have improved the quality of acquired data; however, deconvolutions of the raw MS data to elucidate PGN structures remains a painstaking manual task, where one needs to come up with the potential structures of PGNs (i.e., search space, which can be as large as >6000 structures on ChemDraw)(22, 23) and look for matches of the expected *m/z* values (within a tolerable mass range, usually <5 ppm) in the acquired LC-MS dataset. Recently, Patel *et al.* developed PGFinder, an open-access software that supports the automatic generation of an *in silico* PGN database, which represents a large and well-defined search space for unbiased PGN search.(24) However, for accurate validation of the PGN structure, interpretations of the experimental MS/MS spectra remain a manual task, where one then needs to closely evaluate the MS/MS fragmentation spectra to decipher the particular bond arrangement and connection for each PGN candidate. Such manual annotations of MS data are considerably time-consuming, laborious, and inconsistent, remaining as an undesirable bottleneck for robust and comprehensive peptidoglycan analysis with higher throughput.(25, 26) Manual curation of the PGN library entails drawing the structures of individual PGN molecules to infer their chemical formulae and *m/z* values, which is time-consuming, inconsistent among users, and unlikely to comprehensively enlist all possible structural modifications of peptidoglycans.(22–24) In particular, the manually curated search space and peak assignment in the current workflow are likely inexhaustive since *one only finds what one looks for, if not less*. This may deter the discovery of novel structural features of peptidoglycan, especially in the gut microbiota, where the scope of peptidoglycan diversity has not been much explored (**Figure 1D, top row**). In bacterial peptidoglycan profiling, a targeted search strategy is routinely used for the identification of potential PGNs, where one needs an *a priori* idea of the expected PGN structures to search for the theoretical *m/z* values from the acquired LC-MS dataset.(26)

Towards these challenges, we present a novel and customizable *in silico* PGN MS library to enable automated MS/MS deconvolution for PGN identification (**Figure 1D, bottom row**). Our PGN MS library (.msp format) encompasses the *in silico* predicted MS/MS fragmentation for each PGN in the dataset, which is compatible with open-access and vendor software for automated matching and scoring of the experimental MS/MS peaks, thus streamlining PGN analysis with unmatched confidence and throughput. As our MS/MS prediction algorithm was explicitly developed for PGNs, whose structures embody unique carbohydrate moieties and non-proteogenic amino acids, our PGN analysis pipeline outperforms other existing fragmentation prediction tools in metabolomics and proteomics for PGN identification. For efficient organization of the generated muropeptides, we developed a standardized and descriptive nomenclature for muropeptides (**Figure S1**). Notably, the *in silico* PGN database can be customized according to user-defined parameters (i.e., summarized instructions can be found in **Figure S2A** and a step-by-step guide on database creation is included, **SI1**) and is available for users to download and modify.

Applying the automated PGN analysis pipeline, we profiled the peptidoglycan compositions of five different gut bacteria. Intriguingly, an unusually high abundance of anhydro-PGNs (i.e., PGNs containing a 1,6-anhydro-muramyl moiety (anNAM)) (**Figure 1A, far right**) was found in *Bifidobacterium*. the common probiotic bacteria that confer anti-inflammatory effects in hosts.(27, 28) We further demonstrated that MltG and RfpB homologs in *Bifidobacterium* possess robust lytic transglycosylase (LT) activity towards distinct peptidoglycan substrates to yield anhydro-PGN products. Interestingly, such anhydro-PGNs exhibit potent anti-inflammatory effects *in vitro*, which may open up further opportunities for postbiotic development. Overall, we anticipate that our PGN analysis pipeline will greatly facilitate efforts to explore the structures and functions of gut microbiota-derived PGNs.

## Results

### Generation of a customizable PGN MS1 database

To streamline the PGN searching process, we envisioned a method to automatically generate a PGN MS1 database from user-defined parameters. The basic muropeptide scaffold of PGNs (upon muramidase digestion in the sample preparation) features a (NAG)(NAM) disaccharide with a stem peptide, where distinct structural modifications are possible at each position.(1) To build the PGN database, the user, though a graphical user interface, conveniently selects the possible range of modifications on the (NAG)(NAM) backbone, including *O*-acetylation, de-*N*-acetylation, or anNAM termini, followed by selecting the possible amino acid identities at each stem peptide position. Next, additional structural modifications can be included, such as Braun’s lipoprotein attachment, substitution of the terminal amino acid with lactate in the stem peptide, endopeptidase-cleaved products, and reduction of muramyl termini. Lastly, the user can select the amount and types of PGN polymerization, either through peptide crosslinks or glycosidic bonds. All parameters can be adjusted. Once the parameters are confirmed, PGN molecules are constructed *in silico* with RDKit,(29) and the database is saved as an Excel worksheet (.xlsx). With the graphical user interface, no coding experience from the user is required to build the *in silico* PGN database (**Figure S2** and **SI1**).

Apart from the unique descriptive name for each PGN (**Figure S1**), the *in silico* database (.xlsx) also includes chemical descriptors for individual PGNs, e.g. chemical formula, adducts *m/z*, clogP, InChIKey, SMILES, and PGN-specific descriptors, e.g. the degree of acetylation, degree of amidation, and stem peptide length, thereby facilitating subsequent PGN categorization and comparative analysis (**Figure 2, right**). Accompanying the *in silico* PGN database, an image output that summarizes the user-defined parameters is automatically generated for convenient referencing (**Figure 2, left**). For a typical *in silico* database of 3,000 – 10,000 PGNs, it takes ∼1 min per 1000 PGN to generate when run on a computer with a 2.60 GHz processor and 16 GB RAM.

**Figure 2.**
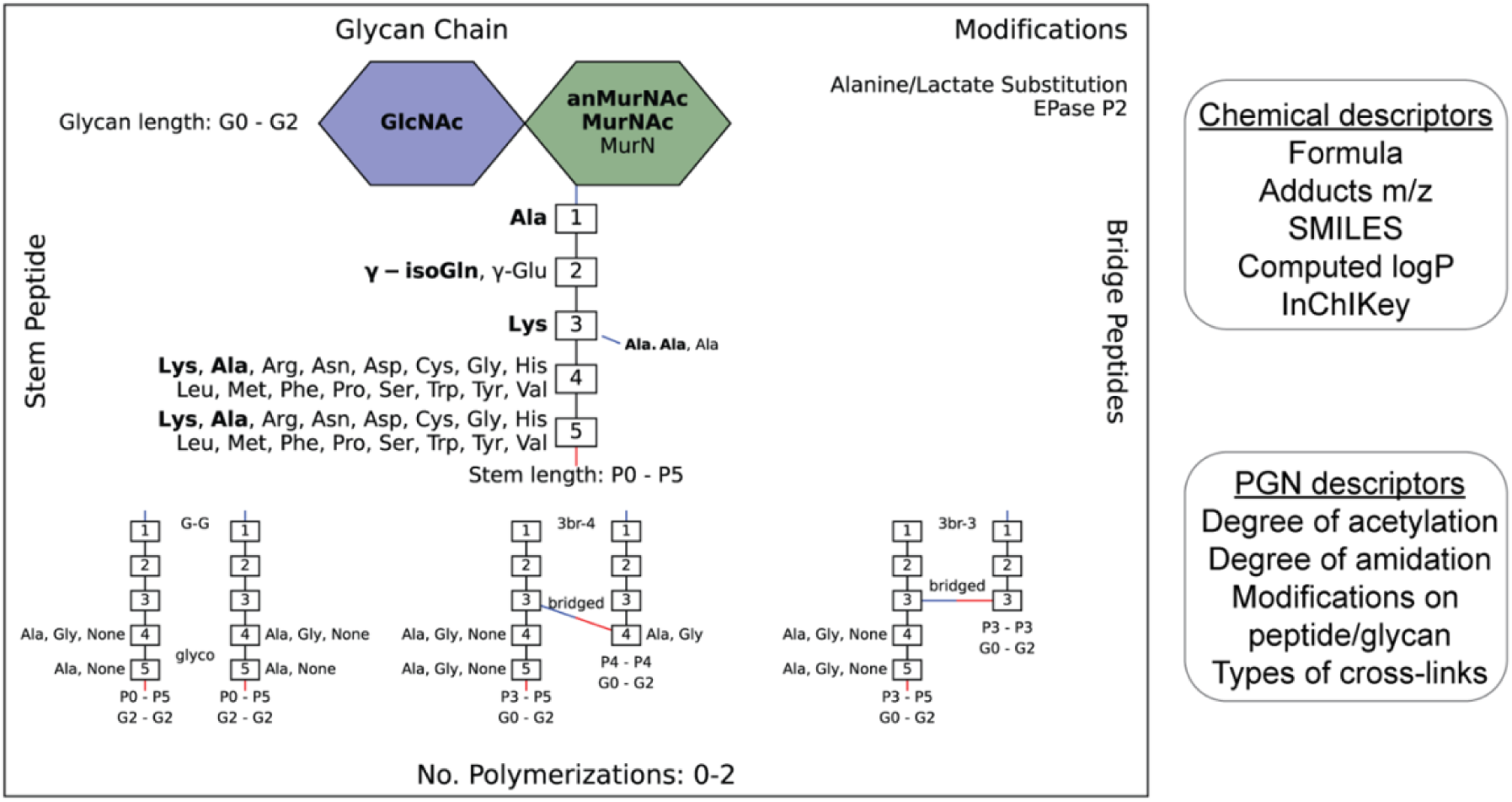
Example of an image output that summarizes the diversity of PGNs in the user-defined *in silico* PGN library (**Left**). The glucosamine (blue hexagon), muramic acid (green hexagon), and amino acids (numbered white boxes) used to construct PGNs are listed. Bridge peptide sequences are indicated to the right of the connecting amino acid with a connecting line. The color of this line indicates how the bridge peptide is connected (red: through the COOH group; blue: through the NH_2_ group.) In creating the *in silico* library, the user may define canonical makeups of peptidoglycans, such as glycans (GlcNAc, MurNAc, anMurNAc), amino acids (Ala, γ-Glu, Lys), and bridge peptides (Ala.Ala), which are shown as bold. The possible glycan chain lengths (0 glycans to 2 glycans), peptide chain lengths (0 amino acids – 5 amino acids), and number of polymerizations (0 to 2 polymerizations) are indicated. Modifications applied are listed in the top-right corner of the image (e.g., alanine/lactate modification and endopeptidase acting on the 2^nd^ stem peptide bond). The requirement of each polymerization is shown at the bottom. For instance, the G-G polymerization is only formed between PGNs with glycan length 2, peptide lengths 0-5, and with either Ala, Gly or no amino acid in positions 4/5 on both acceptor/donor peptidoglycans. All N- and C- peptide termini are colored blue and red, respectively. The list of PGNs in the library is saved in an Excel file (.xlsx) which contains their chemical descriptors (SMILES, InChIKey, chemical formula, etc.) and PGN-specific descriptors (degree amidation, degree acetylation, etc.) (**Right**).

Although an exhaustive PGN search space aids the discovery of low-abundance target molecules, including excessive numbers of structural modifications in monomeric PGNs would result in a combinatorial explosion of the possible PGN polymers. This can significantly compromise data analysis time. Therefore, depending on the potential applications, it may be useful to reduce the number of PGN polymers with a more targeted and focused PGN library. For instance, we recognized that the canonical makeup of *E. faecalis* PGNs contains NAG, NAM, Ala, γ-isoGln, Lys, and Ala-Ala (branch peptide), 0 or 2 glycans (glycan length), 4-5 amino acids (stem peptide length), which altogether account for ∼70% of its total PGNs (**Figure 2, in bold**); hence for building a focused PGN polymer pool, we excluded minor PGN monomers that differ from the canonical makeup (default: >1 difference). In addition, for 3-4 crosslinked PGNs in *E. faecalis,* we also specified the donor and acceptor stem peptides can only include either the canonical (Ala) or the most common substituted amino acid (Gly) at the terminus (**Figure 2, bottom**). By applying such restrictions, the pool of *in silico E. faecalis* PGN dimers generated is significantly reduced from 46,164 to 2,880 (**Table S1**). The species- and crosslink-specific criteria are useful for building a focused *in silico* PGN polymer pool.

### Development of PGN_MS2 for *in silico* MS/MS prediction

Although PGNs can be identified by their *m/z* values alone (MS1 identification), additional analysis by tandem mass spectrometry (MS/MS) is necessary to resolve structural isomers of PGNs with mass coincidences. In the fields of metabolomics and proteomics, compound identification is routinely performed by matching and scoring experimental MS/MS spectra against a reference library of actual MS/MS spectra of standard compounds and/or the *in silico* simulated MS/MS spectra for compounds whose experimental data are not available.(30–33) Given the limited availability of empirically collected MS/MS data, *in silico* MS/MS prediction can greatly improve compound identification.(34) However, due to the unique sugar and amino acid composition present in PGNs, existing MS/MS simulation tools in metabolomics and proteomics are not well-suited for PGN identification.(25) Toward the automated PGN analysis workflow, we next sought to augment the PGN MS1 database with *in silico* predicted MS/MS spectra.

To derive *in silico* PGN MS/MS spectra, we first studied the ESI-MS/MS spectra of known PGNs. Recent studies by Tan *et al.* and Anderson *et al.* reported the experimental MS/MS spectra for selected PGNs from *E. coli*, *S. aureus,* and *P. aeruginosa*, providing a suitable starting point for our evaluation.(22, 35) In addition, we also acquired experimental LC-HRMS/MS data for several major PGNs with known structures from *E. faecalis* and *L. plantarum*. Notably, these spectra were acquired by different MS instruments: namely, Orbitrap Exploris 120 (our study), LCQ Fleet (Tan *et al*), and Q-TOF(Anderson *et al*), which allowed us to derive common ESI-MS/MS fragmentation rules for most PGNs. We recognized that the PGN precursor ions frequently undergo B/Z-type glycan fragmentation and b/y-type peptide fragmentation, with multiple simultaneous b/y cleavages to yield lighter ions (**Figure 3A**). Additionally, the lactoyl bond connecting the glycan and peptide in PGNs also fragments readily, with the peptide fragment ion henceforth named L (**Figure 3A**). Furthermore, isomeric PGNs that contain stem peptides such as Aqm and Aem(NH_2_) with differing amidation positions can be easily distinguished by their MS/MS patterns (**Figure S3**). The y2 peptide fragments (*i.e.* qm or em(NH_2_), *m/z*: 319.1619) undergo further e1/e2 or q1/q2 fragmentations due to prominent neutral losses at the *N*-terminus.(36) For instance, em(NH_2_) yields 301.1465 (e1) and 256.1280 (e2) fragments, whereas qm gives rise to signature MS/MS peaks of 302.1347 (q1) and 257.1103 (q2); with q2 fragments showing higher relative intensities (**Figure S3A-B**). These abundant MS/MS features are useful to distinguish PGNs that bear e or q in the stem peptides, as in the case of *L. plantarum* (**Figure S3**). Upon evaluating the experimental MS/MS spectra for ∼30 PGNs, we found that most of the fragmentation peaks can be explained by 19 fragmentation reactions or a combination thereof (**Figure 3A**).

**Figure 3.**
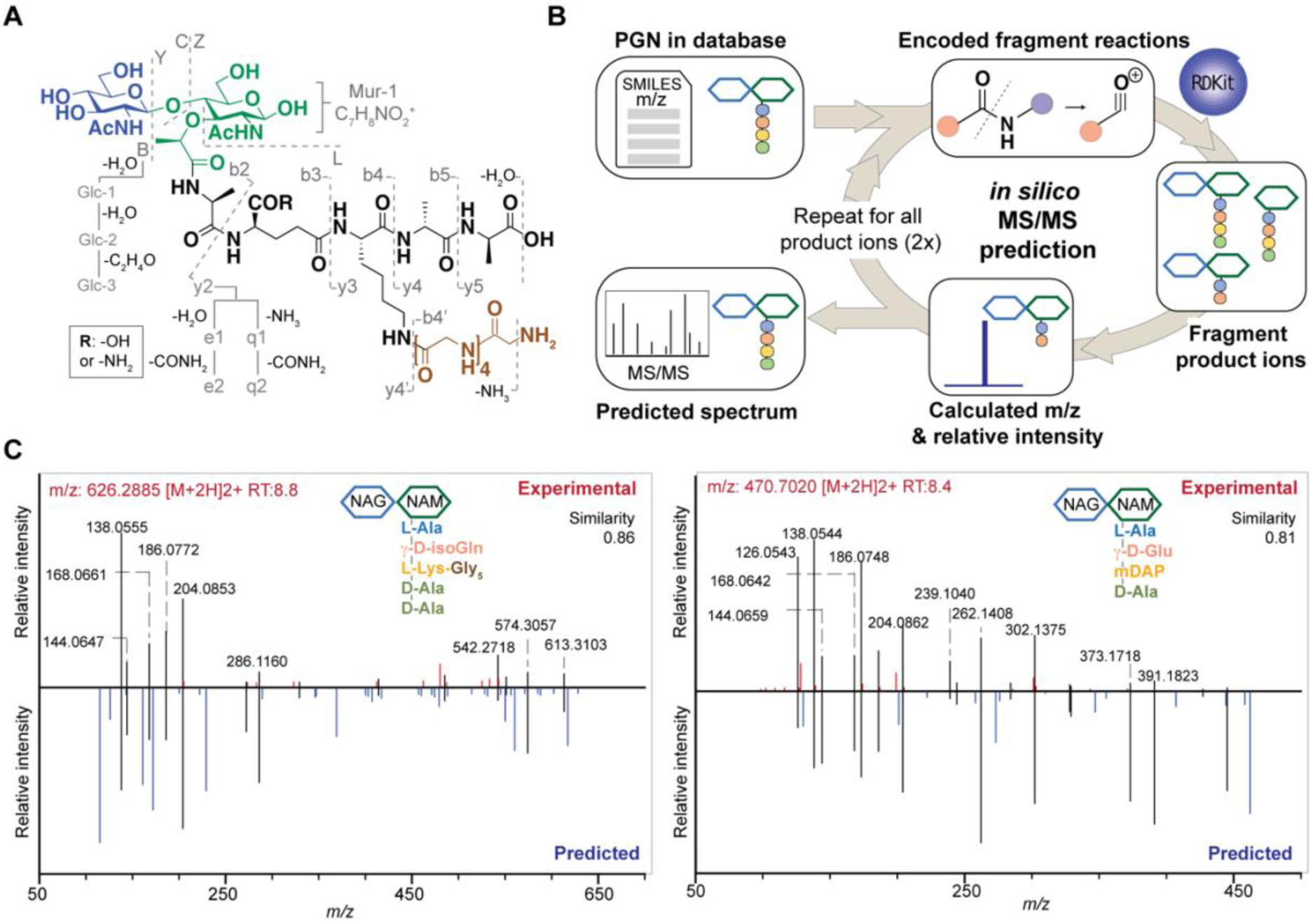
The design and construction of *in silico* PGN fragmenter, PGN_MS2. (**A**) Empirical analysis of PGN MS/MS spectra reveals the possible types of fragmentation reactions, which are encoded in PGN_MS2. Fragmentation of the glycan backbone (B/Y/C/Z) follows the nomenclature from Domon and Costello;(102) fragmentation of the stem peptide is denoted by bn and yn, where n indicates the position of the peptide bond, with the peptide bond nearest to the glycan backbone denoted as “1”; fragmentation of the bridge peptide is indicated with a quotation mark (’). In addition, the γ-Glu-containing PGNs yield e1/e2 fragments due to the neutral loss of H_2_O and COOH+NH_3_, respectively; similarly, γ-isoGln-containing PGNs generate q1/q2 fragments by neutral loss of NH_3_ and CONH_2_+NH_3_, respectively. Furthermore, further fragmentation of GlcNAc / MurNAc (Glc-1, Glc-2, Glc-3, Mur-1) with neutral loss of H_2_O or NH_3_ are also included as fragmentation reactions. **(B)** The schematic of *in silico* MS/MS spectrum generation. The fragmentation reactions for each PGN are encoded as SMARTS, where the *m/z* and relative intensity is calculated for each product ion. Each product ion undergoes further fragmentation (2 repeats) to create the *in silico* spectrum. **(C)** Comparison of experimental MS/MS spectra (top, red) vs. *in silico* predicted spectra (bottom, blue) for canonical PGN in *E. coli* (left) and *S. aureus* (right). Matched peaks are coloured black, and the cosine similarity scores between the spectra are shown in the top right corner.

Based on these common fragmentation reactions, we developed the PGN_MS2, an *in silico* MS/MS prediction tool for PGNs. As shown in **Figure 3B**, we encoded each fragmentation as a chemical reaction in SMARTS and simulated it with RDKit.(29) Each parental PGN ion (generation-0) is fragmented via all possible 19 reactions to form generation-1 product ions, which are further fragmented to yield generation-2 and generation-3 product ions sequentially. Fragmentation is discontinued after no new product ions are generated. For every fragmentation, the *m/z* value and relative intensity for each fragment are calculated. Relative intensity is estimated based on an empirical formula (that accounts for the number of peptide bonds or mass ratio of the precursor and product ions) together with a fragmentation-specific adjustment factor. Finally, the assembly of possible fragment ions affords the *in silico* predicted MS/MS spectra. To account for the different precursor adducts (i.e., [M+H]^+^, [M+2H]^2+^, and [M+3H]^3+^), separate MS/MS spectra are created for each adduct, whereby fragment ions with *m/z* greater than the precursor ion are removed. In sum, our PGN library integrates the predicted MS/MS spectra of all *in silico* PGNs in the database as a NIST format text file (.msp, **Figure S2C**).

### Reliable PGN identification with *in silico* MS/MS prediction

To test the accuracy and reliability of MS/MS prediction by PGN_MS2, we first compared the experimental spectra of a panel of distinct PGNs from different bacteria with their respective predicted spectra by calculating the cosine spectral similarity scores.(37) As expected, PGN_MS2 consistently afforded high similarity scores of 0.7-0.8 for most PGNs, which significantly outperformed other spectral prediction tools such as CFM-ID and ms2pip (**Figure 3C and S4, Table S2**),(32, 33) showcasing the specialized applications of PGN_MS2 for PGN analysis.

Next, we confirmed that the *in silico* predicted spectra by PGN_MS2 match well with the MS/MS spectra acquired by either an Orbitrap spectrometer via higher-energy C-trap dissociation (HCD)- based fragmentation or a Q-TOF instrument via collisional dissociation (CID)-based fragmentation (**Figure S5A-D**).(38) In addition, to benchmark our PGN_MS2 with the available PGN dataset, we also evaluated the experimental data of *P. aeruginosa* PGNs deposited by Anderson *et al*, which was collected by a Q-TOF mass spectrometer.(22) Consistently, we readily confirmed 58 out of 62 of the PGNs that were manually identified in the previous work (**Figure S5E-F**). Taken together, our observations demonstrate the robustness and reliability of PGN_MS2 in simulating ESI-MS/MS spectra of PGNs for structural determination.

To further interrogate if PGN_MS2 could indeed aid accurate assignment of PGNs among closely related structural isomers, we challenged it to identify the canonical *E. coli* or *S. aureus* PGN, (NAG)(NAM)-AemA and (NAG)(NAM)-AqKAA[3-NH2-GGGGG] respectively, from a set of four intentionally generated mock PGNs with identical molecular formulae (**Figure S6**). Satisfactorily, we correctly assigned the two PGN structures, since they both emerged as the top hits with the highest spectral similarity scores compared to other possible isomers, albeit to a small margin (**Figure S6**). Based on our analysis, we noted that although the top matched *in silico* PGN usually represents the accurate structure, other criteria such as the presence or absence of certain signature MS/MS fragments are particularly useful for PGN determination too. For instance, fragments containing the intact mDAP-mDAP bond (*i.e.*, *m/z*: 617.2777, 746.3203, 889.3785) are observed in the MS/MS spectra of the 3-3 but not 3-4 crosslinked PGNs in *E. coli*, allowing convenient distinction between the two isomers (**Figure S4C and Figure S7**). Therefore, it is prudent to check for these signature fragments for PGN identification. To assist with this, PGN_MS2 also annotates the chemical structures of each fragment in the predicted MS/MS spectra as SMILES (**Figure S2D**).

### Validation of the MS/MS-integrated workflow for model bacterial PGN profiling

Upon demonstrating the reliability of PGN_MS2 for identifying individual PGN molecules, we then sought to evaluate its potential application for profiling bacterial peptidoglycan compositions. We first constructed *in silico* PGN MS libraries customized for different model bacteria, including *E. coli*, *S. aureus*, *E. faecium*, and *L. plantarum*. We next acquired experimental LC-HRMS/MS data of PGNs from these bacteria. NaBH_4_ reduction was omitted to prevent potential acid hydrolysis during the addition of phosphoric acid and preserve the natural structure of PGN. Next, we utilized open-source software MS-DIAL for automated data analysis by importing the respective *in silico* MS/MS libraries as spectral references for PGN identification.(39) In general, our findings are consistent with previous knowledge of PGN compositions in these bacteria,(21, 24, 40–47) validating our MS/MS-integrated PGN MS library for automated PGN profiling. We summarized the canonical PGN monomeric makeup (**Figure 4A**) and listed the detailed PGN compositions in these bacteria (**Table S3-8**). Below we highlight the discovery of several PGN structural features that exemplify the virtue of the *in silico* MS/MS spectral library.

**Figure 4.**
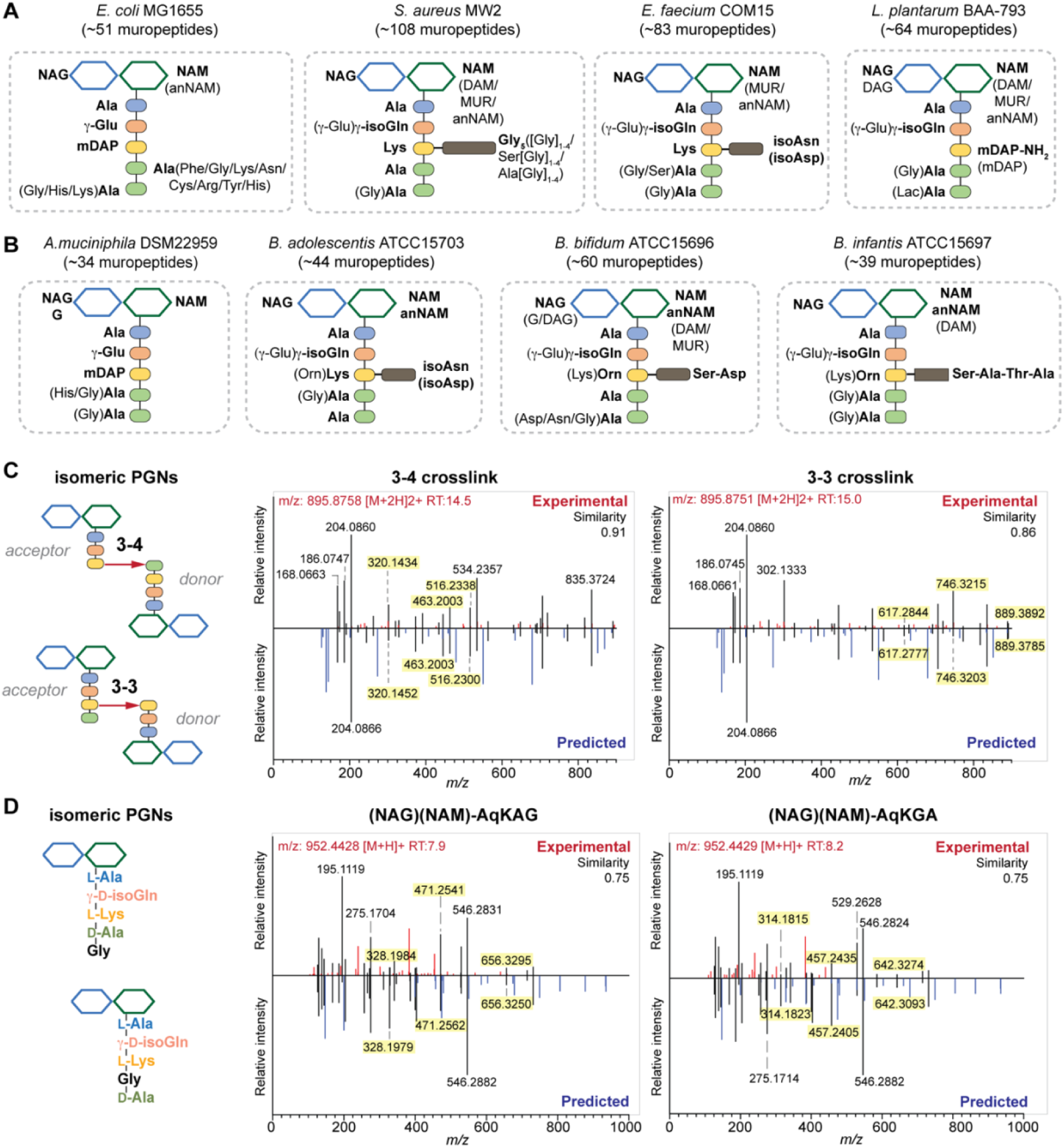
Summary of peptidoglycan compositions in the model (**A**) and gut bacteria (**B**) with the canonical makeup shown in bold and variable components not bolded. For instance, the canonical peptidoglycan makeup in *E. coli* is (NAG)(NAM)-AemAA and the fifth amino acid, Ala, can be substituted with His, Gly, or Lys. The total number of PGNs identified in each bacteria is listed. (**C-D**) PGN_MS2 enables distinctions of isomeric PGNs. In *E. coli*, the tetrapeptide-tripeptide dimers with either 3-4 or 3-3 crosslinks are resolved by matching experimental spectra with *in silico* predicted MS/MS patterns (**C**). In *E. faecium*, monomeric PGNs with Gly either at the 4^th^ or 5^th^ position in stem peptide. The isomeric PGNs yield distinct MS/MS fragments that can be distinguished with PGN_MS2 (**D**). The key fragments that are essential for resolving respective isomers are highlighted in yellow. **Figures S7-8** showcase additional examples of differentiating isomeric PGNs by PGN_MS2.

Amidation of the stem peptides is a unique feature in PGNs of Gram-positive bacteria (**Figure 5B**).(1) For instance, the canonical monomeric PGNs in *E. faecium* and *L. plantarum* each contain two possible amidated residues in the stem peptides, q and isoAsn, q and m(NH_2_), respectively (**Figure 4A**). Although most PGNs in both bacteria are amidated at both positions, substantial amounts of the singly amidated PGNs are also observed, which require MS/MS analysis to determine the exact amidation position in the isomeric PGNs (**Figure S3**). In addition, some *L. plantarum* PGNs have D-lactate instead of D-Ala at the peptide terminus,(47) which further complicates identification. The three structural isomers, (NAG)(NAM)-Aem(NH2)AA, (NAG)(NAM)-AqmAA, and (NAG)(NAM)-Aqm(NH2)ALac have identical *m/z* values that are indistinguishable solely based on MS1 analysis and require in-depth MS/MS evaluation. With our approach, the *in silico* predicted MS/MS spectra by PGN_MS2 revealed signature fragments for each of the three PGN isomers, which significantly improve the confidence and throughput of MS/MS identification (**Figure S8**). For instance, the experimental spectra of (NAG)(NAM)-Aqm(NH2)Alac showed the best match to the *in silico* spectra for this particular isomer and contains all key fragments, allowing us to easily assign the correct structure (**Figure S8**). Moreover, with our MS/MS-integrated analysis pipeline, we also uncovered that amidation at the second residue (q) of the stem peptide is more prominent than at the side chain (β-Asp) in *E. faecium*, whereas similar amidation rates were observed for both q and m(NH_2_) in PGNs of *L. plantarum* (**Figure S10A**).(47) Recognizing that bacterial peptidoglycan amidations are associated with increased levels of crosslinking and also implicate antibiotic resistance,(48–53) we anticipate that our workflow for the facile analysis of such amidated PGNs will facilitate the development of novel antimicrobials targeting bacterial peptidoglycan amidations.

**Figure 5.**
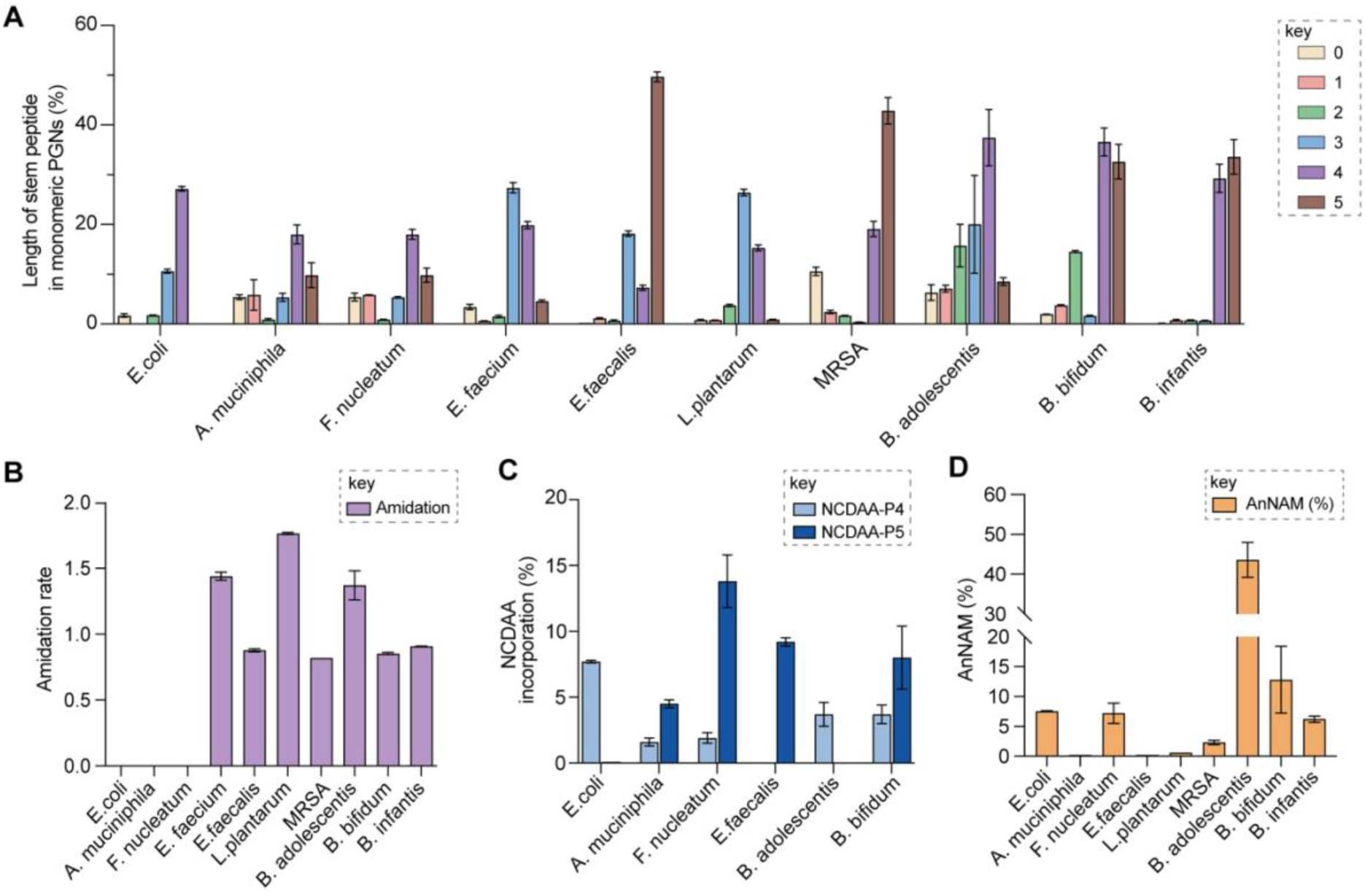
Summary of peptidoglycan features in model and gut bacteria. (**A**) Profiling of monomeric PGNs with varying numbers of amino acids in the stem peptide across bacteria. (**B**) Profiling of amidation status in stem peptides across bacteria. (**C**) Profiling of NCDAA incorporations in stem peptides across bacteria. (**D**) Profiling of the amount of anNAM termini in bacteria, where *Bifidobacterium* spp. showcase a high abundance of anNAM that differs from typical Gram-positive bacteria. All statistics indicate the relative muropeptide composition (in %) except for (**B**), where the amidation rate is defined as the frequency of amidated residues (γ-D-isoGln/β-D-isoAsn/mDAP(NH2)) per muropeptide. *L. plantarum*, *E. faecium*, and *B. adolescentis* feature two amidated amino acids, and the values shown are the combined rates for both. The data represent the average of three to four biological replicates with error bars representing standard deviations. Additional profiling analysis is in **Figure S10**.

Peptidoglycan crosslinking via stem peptides confers strength and resistance to certain antibiotics and stress conditions. For instance, *E. coli* typically manifests 3-4 crosslinking but significantly increases 3-3 crosslinking during stress conditions.(54, 55) The 3-4 and 3-3 crosslinked tripeptide-tetrapeptide dimeric PGNs are structural isomers that differ only in the isopeptide bond position, which were easily distinguished using our MS/MS-integrated PGN analysis workflow (**Figure S4C and S7**). Interestingly, across all bacteria, we also detected tetra-saccharide PGN dimers that are isomeric to the crosslinked dimers (**Figure S11**). Although such tetra-saccharide motifs are possible products of incomplete muramidase digestion during sample preparation, additional rounds of enzymatic digestions could not fully eliminate them.(56) Compared to the crosslinked PGN dimers, these tetra-saccharide PGNs generally yielded fewer MS/MS fragments with lower relative intensity for B-type fragments and higher intensity for L-type fragments (**Figure S11**), which is consistent with the presence of only one terminal GlcNAc and two free-stem peptides in these structures. The ability to easily identify such tetra-saccharide PGNs in our workflow may provide the impetus to investigate their physiological relevance in bacteria.

Notably, *S. aureus* is well-recognized for possessing a complex peptidoglycan structure due to its extended bridge peptide and a multitude of PG remodelling enzymes. Previous studies by HPLC UV quantification methods revealed the highly crosslinked (crosslinking index ∼90%) peptidoglycan in *S. aureus*.(57) However, in our LC-MS analysis of the soluble muropeptide fraction for *S. aureus*, we were unable to detect any oligomers larger than trimers (**Table S8**). These oligomers would form anomers (arising from mutarotation of C_1_-OH on the muramyl sugar)(20) and hence present as multiple, smaller peaks below the limit of detection, owing to the poor ionizability of the larger oligomeric species. Therefore, we noted that the percentage of oligomers is diminished in our analysis, yielding a crosslinking index of only ∼8%. Nevertheless, a diverse range of PGNs in *S. aureus* were annotated (149 muropeptides, **Table S8**). For instance, apart from the canonical pentaGly-containing PGNs in *S. aureus* (∼50%, **Table S8**), we also observed a substantial amount of PGNs containing extra Gly residues,(21, 41, 42) which were confirmed to be appended at the termini of the stem peptide via MS/MS analysis (**Figure S11**). These PGNs are likely the cleavage products of glycyl endopeptidases (i.e. lysostaphin-like enzymes LytM or LytU) acting on cross-linked PGNs,(58, 59) which altogether account for ∼40% of the total PGNs, suggesting extensive peptidoglycan remodelling and hydrolysis of *S. aureus* at stationary phase (**Table S8**). Moreover, additional amidase- and glucosaminidase-cleaved PGNs were also detected at ∼10% and ∼1% abundance respectively (**Table S8**). Such heightened peptidoglycan hydrolase and remodelling activities in bacteria could be indicative of peptidoglycan recycling, the release of cell wall-anchored proteins, the construction of secretion systems, or modulation of cell wall elasticity.(60–62)

### Comprehensive and automated PGN profiling in gut bacteria

Encouraged by the proof-of-concept studies in model bacteria, we next set out to comprehensively profile the PGNs in a panel of human gut bacteria: *Akkermansia muciniphila*, *Bifidobacterium adolescentis*, *Bifidobacterium bifidum*, *Bifidobacterium infantis*, and *Fusobacterium nucleatum*. Among them, *A. muciniphila* and the *Bifidobacterium spp*. are commensal species that help to maintain the gut microbiota balance and reduce inflammation, whereas *F. nucleatum* is associated with colorectal and other cancers. (27, 28, 63–65) Notably, except for a recent study that broadly categorized PGNs in *A. muciniphila* using LC-MS,(66) our knowledge of *Bifidobacterium* and *F. nucleatum* PGNs is only from early studies in the 1970s. (67–70) To address their potential biological functions in the host, there is an imperative need to perform in-depth PGN profiling of these gut bacteria.

We first elucidated the canonical PGN makeup in the respective gut bacteria (**Figure 4B**). *A. muciniphila* possesses mDAP-type PGNs,(66) similar to most other Gram-negative bacteria. However, *F. nucleatum* PGNs exclusively feature the non-proteinogenic lanthionine at the third position of the stem peptide, whose structure closely resembles that of mDAP.(68, 69) On the other hand, Gram-positive *Bifidobacterium spp.* possess either L-Lys or L-Orn as the third residue that is further appended with distinct bridge peptides (**Figure 4B**).(67) Surprisingly, we found that whereas the L-Lys containing PGNs are only minor constituents in *B. bifidum* and *B. infantis* (2.7 and 3.6% respectively, **Figure S10B**), they are the dominant ones in *B. adolescentis* (62.3%, **Figure S10B**). This could imply that MurE, the ligase that incorporates the third amino acid residue in soluble peptidoglycan precursors, exhibits unique substrate tolerances amongst different species of *Bifidobacterium.* Furthermore, *B. adolescentis* PGNs also sport an identical bridge peptide (i.e., β- Asp/β-isoAsn) as those in *E. faecium* and *L. lactis,*(46, 71, 72) which are constructed by the sequential enzymatic activities of the D-aspartate ligase, Aslfm, and the asparagine synthase, AsnH (71–73). Consistently, *B. adolescentis* encodes homologs of both enzymes (**Table S14**).

Evaluating the lengths of stem peptides in PGNs across different bacteria, we found that PGNs in *F. nucleatum*, *B. infantis,* and *S. aureus* predominantly possess penta- and tetra-peptides, whereas *B. adolescentis, L. plantarum,* and *E. faecium* showcase variable PGNs with shorter stems ranging from one to four amino acids, which are likely products of enzymatic cleavages by DD-carboxypeptidases, LD-endopeptidases or DL-endopeptidases during PG maturation in bacteria (**Figure 5A**).(19) Recent studies have revealed that SagA-like DL-endopeptidases secreted by commensal gut bacteria such as *E. faecium* and *Lactobacillus* generate bioactive PGN motifs that regulate host gut homeostasis.(46, 74, 75) Interestingly, both *B. adolescentis* and *B. bifidum* show a significant proportion of PGNs with dipeptide stems (∼15%) (**Figure 5A**), suggesting the activities of SagA-like enzymes in these two *Bifidobacterium* that could be potentially relevant to their anti-inflammatory effects.

In all bacteria, NCDAAs are commonly found in the stem peptides of PGNs, substituting D-Ala in the fourth or fifth position (**Figure 4A-B** and **5B**).(17) PGNs from *A. muciniphila* and *F. nucleatum* mostly contain basic NCDAAs such His, Arg, Asn, or Lys at the fifth position of the stem peptides (**Figure 4B**), which could be incorporated by transpeptidases and/or Ddl in these bacteria. (76) Notably, *E. coli* possesses the greatest diversity of NCDAAs in PGNs, including Phe, Tyr, Gly, Lys, Cys, Arg, etc., whereas other bacteria, *B. adolescentis*, *B. infantis*, *L. plantarum,* and *S. aureus* appear to solely utilize Gly as the non-canonical amino acid in the PGN stem peptides (**Figure 4A-B**). Empowered by *in silico* MS/MS spectral references, we readily distinguished PGN isomers with Gly at either the fourth or fifth position of the pentapeptide stem in *E. faecium* (**Figure 4D**). NCDAAs in peptidoglycan confer bacterial resistance against hydrolases of rival bacterial species, which are consistently found at elevated levels in bacteria under stress conditions.(17, 77) Our work reveals the widespread presence of NCDAAs in bacterial PGNs at steady-state conditions than previously appreciated.

Besides stem peptide motifs, we also profiled structural features on the (NAG)(NAM) backbone in PGNs across bacteria, including *O-*acetylation (i.e., DAG, DAM) or de*-N-*acetylation (i.e., G, MUR) (**Figure 4A-B**).(78) Modifications to acetylation in peptidoglycan may help bacteria evade lytic enzymes such as lysozyme.(4) With our MS/MS-integrated analysis workflow, we could readily determine if acetylation/de-acetylation occurs on the NAG or NAM residue in the disaccharide PGNs. Such alterations only account for a minor extent (<5%) in PGNs of *B. bifidum* and *L. plantarum*, hence no significant changes in the overall acetylation rate of PGNs were observed for most bacteria (**Figure S10E-F**). One remarkable exception is *A. muciniphila* that showcases 43% de-*N-*acetylation of NAG (**Figure S10E-F**), which is in good agreement with the recent analysis by Garcia-Vello *et al.* (∼40%).(66) Notably, these de-*N*-acetylated PGNs are still potent agonists to both NOD1 and NOD2 immune sensors;(66) thus, it remains to be determined if such de-*N*-acetylated motifs exhibit any distinct functions in the host.

Next, 1,6-anhydroMurNAc (anNAM) termini are unique features that mark the end of the peptidoglycan strands in Gram-negative bacteria.(26) Correspondingly, anhydro-PGNs constituted 4 – 5% of total peptidoglycan composition in Gram-negative bacteria, *E. coli* and *F. nucleatum*, but are nearly undetectable in model Gram-positive bacteria and *A. muciniphila* (**Figure 5D**).(66) Surprisingly, we found that all three *Bifidobacterium spp*. contain a remarkably high abundance of anhydro-PGNs, which is unusual for Gram-positive bacteria (**Figure 5D and Figure S9**). For instance, the anNAM-containing PGNs comprise nearly 40% of total PGNs in *B. adolescentis* (**Figure 5D**). The exceedingly high amounts of anhydro-PGNs in *Bifidobacterium* suggest the presence of highly active lytic transglycosylases (LTs) in catalyzing the non-hydrolytic cleavage of peptidoglycan backbone, which are elusive in Gram-positive bacteria.(56, 79, 80) We next set out to establish the putative LTs in *Bifidobacterium* responsible for anhydro-PGNs formation.

### Identification and characterization of the putative LTs in *Bifidobacterium*

To identify putative LTs in *Bifidobacterium*, we searched for homologous proteins containing the catalytic domains of known LTs in other species (i.e. *E. coli* MltA-G, Slt70)(81) on UniProt.(82) Interestingly, all three *Bifidobacterium* species encode proteins (BaMltG, BbMltG and BiMltG) that possess the catalytic domain, IPR003770, of MltG, consistent with the broad conservation of MltG across bacteria (**Figure S13**).(83) Protein sequence alignment with ClustalOmega revealed that BaMltG, BbMltG, and BiMltG are ∼55-57% similar to one another, and are 23-27% similar to *E. coli* MltG, *B. subtilis* MltG, and *S. pneumoniae* MpgA, whose biochemical activities have been characterized (**Figure S14**).(56, 83, 84) While MltG in Gram-negative bacteria represents the sole inner membrane-bound LT that is responsible for the cleavage of the nascent PG strands and generates 1,6-anhydro-MurNAc termini, the MltG homolog in *S. pneumoniae*, MpgA acts as a muramidase instead.(56) Notably, the identity of a single amino acid in the active site of MltG serves as a key determinant for the corresponding enzymatic activity, with Asp for LTs and Asn for muramidases.(56) We found that *Bifidobacterium* MltG harbours an Asp at this position, implying its potential role as an LT (**Figure S14**). For biochemical characterization of *Bifidobacterium* MltG, we cloned, overexpressed, and purified the respective MltG lacking the *N*-terminal transmembrane domain (**Figure 6B**). Initial attempts of incubating the recombinant MltG protein with bacterial sacculi did not yield any products, which were in agreement with previous findings that mature sacculi are not suitable substrates of MltG.(56, 83, 84) We next sought to evaluate the activity of *Bifidobacterium* MltG with nascent peptidoglycan as substrate, which was typically generated from *in situ* Lipid II polymerization.(56, 84) Since it is challenging to isolate native Lipid II molecules from large-scale cultures of *Bifidobacterium*, we used *E. faecalis* Lipid II instead,(85) whose structure closely resembles that of *Bifidobacterium* Lipid II. Upon SgtB polymerization of Lipid II, we added *Bifidobacterium* MltG followed by mutanolysin to release soluble muropeptides for LC-MS analysis. As shown in **Figure 6A-B**, the addition of *Bifidobacterium* MltG indeed led to a significant increase in anhydro-PGN products, indicating its robust LT activity *in vitro* (**Figure 6C and Figure S15**). As a negative control, we showed that mutating the catalytic residue Asp to Ala in *Bifidobacterium* MltG completely abolishes the observed LT activity *in vitro*. (**Figure S16**)

**Figure 6.**
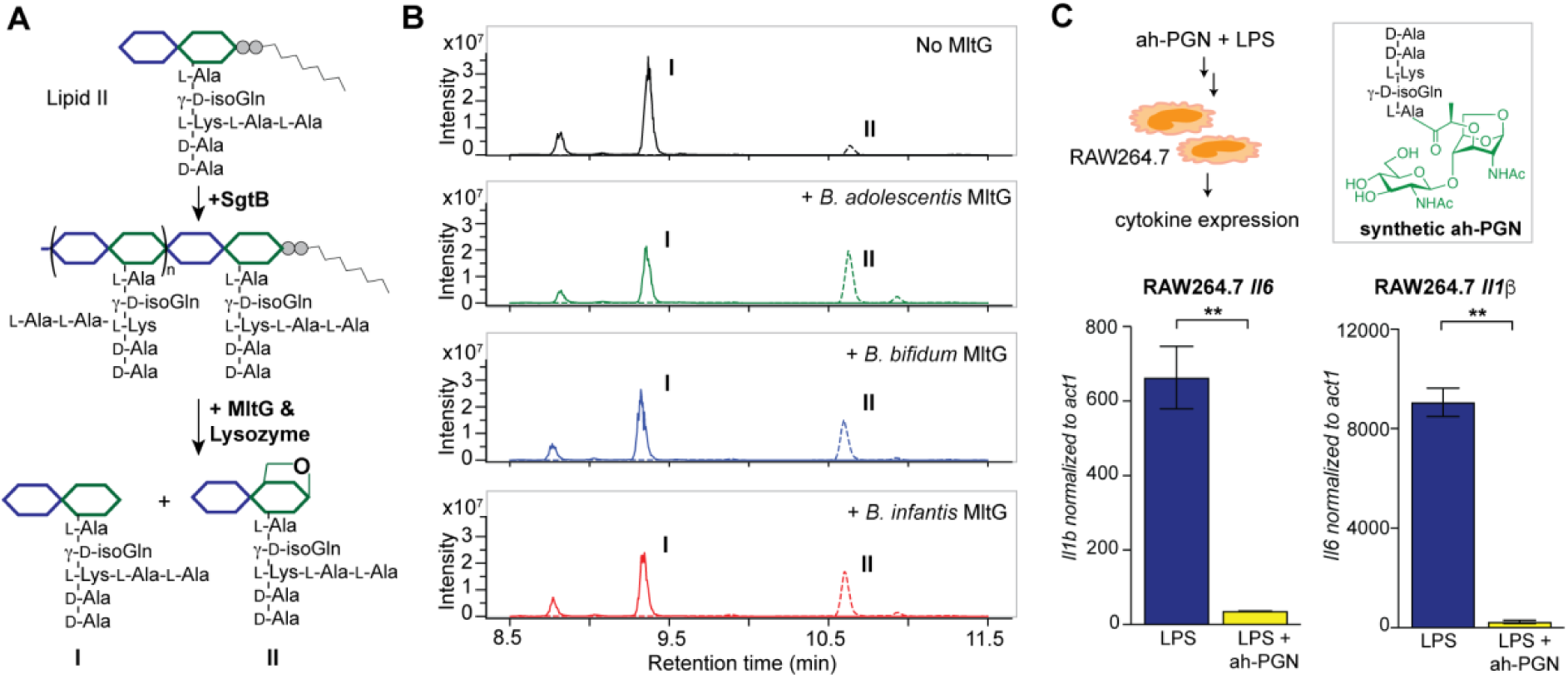
*Bifidobacterium* anhydro-PGNs from the cleavage of lytic transglycosylases (LTs) exhibit potent anti-inflammatory effects *in vitro*. (**A-B**) Biochemical reconstitutions of recombinant *Bifidobacterium* MltG with nascent peptidoglycan as substrates (**A**); LC-MS chromatograms of the muropeptide products indicate the formation of anhydro-PGNs, II (**B**). Extracted ion chromatograms (EICs) for [M+2H]^2+^ were shown. Lipid II was extracted from *E. faecium*. (**C**) Pre-treatment of synthetic anhydro-PGN (ah-PGN), (NAG)(NAM)-AqKAA, significantly suppresses LPS-induced inflammatory responses in murine macrophage RAW264.7 cells. The synthetic ah-PGN mimics the natural anhydro-PGNs found in *B. adolescentis*.

Apart from the well-characterized LTs in Gram-negative bacteria, certain Gram-positive bacteria that undergo dormancy also encode a large family of cell wall lytic enzymes that are known as resuscitation-promoting factors (Rpfs), some of which are LTs.(86, 87) Since *Bifidobacterium* can also enter viable but non-culturable (VBNC) state similar to dormancy,(88, 89) we explored if *Bifidobacterium* could possess any Rpfs with LT activity. Using sequence similarity searching by BLAST, we identified two candidate proteins (RpfB-FL, RpfB-truncated) in *Bifidobacterium* containing the lysozyme-like domain (IPR023346) that show weak homology to *M. tuberculosis* and *S. coelicolor* RpfB (**Figure S13**). Interestingly, both the full-length and truncated RpfB proteins of *B. adolescentis* display dual LT and amidase activities with bacterial sacculi *in vitro* (**Figure S17**), indicating their possible involvement in sacculi remodelling. Taken together, our results established three *bona fide* LTs (MltG, RpfB-full length, RpfB-truncated) in *Bifidobacterium* that may act in concert contributing to the high abundance of anNAM in *Bifidobacterium* peptidoglycan.

### *Bifidobacterium* anhydro-PGNs exhibit potent anti-inflammatory activity *in vitro*

Intrigued by the predominant anhydro-PGNs in *Bifidobacterium spp.*, we hypothesize that the remarkable anti-inflammatory functions of *Bifidobacterium spp.* as probiotics may be attributed to these unique anhydro-PGN molecules. Although most bacterial PGNs belong to pathogen-associated molecular patterns (PAMPs) that are agonists of the mammalian NOD immune sensors to trigger downstream proinflammatory responses,(90) these anhydro-PGN motifs in *Bifidobacterium spp.* lack critical structural features for both NOD1 and NOD2 activations. Specifically, *Bifidobacterium* PGNs harbour an L-Lys or L-Orn instead of mDAP in the stem peptide, rendering them non-agnostic to NOD1;(91, 92) moreover, these PGNs with 1,6-anhydro-MurNAc termini effectively evade NOD2 recognition, which strictly senses the reducing-end anomeric configuration of MurNAc in PGNs.(93, 94) As expected, we demonstrated that crude PGNs of *B. adolescentis* exhibit significantly reduced capacity in activating the NOD signalling pathways in the cell-based reporter assays compared to PGNs of other Gram-positive and Gram-negative bacteria including *S. aureus*, *E. faecalis,* and *E. coli* (**Figure S18**), highlighting the distinct characteristics of PGNs from probiotic *Bifidobacterium spp*. To further explore the potential anti-inflammatory effects of these anhydro-PGNs, we resorted to an *in vitro* immunological assay, where we pre-treated murine macrophage RAW246.7 cells with a synthetic anhydro-PGN before the addition of LPS, followed by gene expressions analysis by RT-qPCR (**Figure 6D** and **Figure S18**).

To our surprise, the presence of the anhydro-PGN effectively suppressed the expressions of several key proinflammatory cytokines including *tnfα, il1b*, and *il6* in RAW246.7 induced by LPS, highlighting the potent anti-inflammatory properties of *Bifidobacterium* anhydro-PGNs *in vitro*. Since these anhydro-PGNs are not agonists of the canonical NOD signalling pathways, the mechanistic details of their anti-inflammatory activities remain to be determined.

## Discussion

Peptidoglycan, an evolutionarily conserved exoskeletal macromolecule that is present across all bacteria, has been increasingly recognized as a key effector of the gut microbiota in regulating host biology.(13–15) While peptidoglycan is largely a meshwork of NAG-NAM glycopolymers with short peptide crosslinks, the diverse structural modifications on the glycan backbone and variations of the amino acid compositions in peptidoglycans of different bacteria have a profound impact on the bioactivities in the host. Notably, the chemical compositions and structural characteristics of bacterial peptidoglycan have been the subject of intense studies since the 1950s, with pioneering efforts to address the identities of amino acids and sugars in peptidoglycan using two-dimensional paper chromatography.(95) In 1988, Glauner established an HPLC-based workflow to separate and quantify soluble PGNs from muramidase digestion of peptidoglycan polymer, where the individual peaks are then collected and subjected to MS for structural determination.(20) The use of HPLC and MS has revolutionized the field and become the standard protocol for PGN analysis. Over the years, such analytical pursuits have revealed many complex structural features in bacterial peptidoglycan that were previously uncharacterized. The advances in modern HPLC-MS instruments in recent years have further greatly improved the speed, resolution, and sensitivity of PGN data acquisition, yet the MS data analysis remains a highly manual process that impedes the characterization and discovery of novel structural features in PGNs in the gut microbiota. In this work, we developed an open-access, customizable MS/MS-integrated PGN library that enables automated PGN searching and identification with the open-source program MS-DIAL,(39) rendering the entire PGN analysis workflow more accurate, robust, and accessible than ever.

Our MS/MS-integrated PGN library is a versatile collection of PGN molecules with *in silico* predicted MS/MS fragmentation spectra. Centred around the (NAG)(NAM)-containing disaccharide muropeptides as the core structure, which embodies the fundamental chemical makeup of bacterial peptidoglycan polymer, the PGN database is customized with user-defined parameters to accommodate diverse structural modifications and polymerizations/crosslinking in PGNs. Currently, our algorithms support most known PGN modifications as built-in selections, including *O*-acetylation, de-*N*-acetylation, NCDAA incorporation, 3-3/3-4 crosslinking, etc; additional structural features can be conveniently incorporated to expand the search space for identification of novel PGNs in the gut microbiota. For each PGN candidate in our database, an *in silico* predicted MS/MS spectrum is automatically generated, which collectively affords a comprehensive *in silico* PGN spectral library. Although the “gold standard” of compound identification by LC-MS/MS is to compare the experimental MS/MS spectra with authentic standards of known structures,(34) the limited availability of empirical MS data for PGNs renders this impractical, since PGNs are not covered in any existing metabolite libraries. In the fields of metabolomics and proteomics, MS spectral prediction algorithms such as LipidBlast, CFD-ID, and ms2pip, etc are widely used to simulate *in silico* MS/MS fragmentation for molecules based on rules (derived manually or with machine learning);(30–33) however, these tools are not well-suited for the PGN chemotypes.(25) To derive specific ESI-MS/MS fragmentation rules for PGNs, we evaluated the experimental MS/MS spectra for ∼30 PGNs of different bacteria acquired by different MS instruments and identified 19 common fragmentation reactions for PGNs. We then developed the PGN_MS2 tool that encodes these *in silico* reactions to simulate MS/MS spectra of PGNs in the MS1 database and demonstrated the use of the open-source software MS-DIAL for data processing, which easily performs the searching and scoring of the experimental data against our *in silico* PGN spectral references. Compared to PGFinder,(24) a recent novel PGN analysis pipeline based solely on MS1 values, our PGN MS library integrates the *in silico* MS/MS spectral prediction for more robust and accurate identification of PGN structural isomers in an automated analysis workflow. Similar to the iterative searching strategy in PGFinder,(24) we also recommend users specify selective parameters to build the *in silico* PGN polymer pool focusing on the major canonical features in the PGN monomers, thereby reducing the combinatorial explosion of the possible polymers. As part of our PGN library, an image summarizing the user-defined parameters in building the *in silico* PGN database, as well as the list of PGN nomenclatures with their respective chemical- and PGN-specific properties are automatically generated as output, illustrating novel features of our workflow.

Notably, our PGN_MS2 represents a dedicated *in silico* MS/MS spectral prediction algorithm for PGNs, whose unique sugar and non-proteogenic amino acids defy reliable predictions by the existing tools developed for small molecules or proteins. Although specialized proteomics analysis software such as Byonic has been applied for PGN analysis,(25) it utilizes a peptide-centric approach, where common PGN structures are viewed as variable modifications of the stem peptide. For instance, one needs to manually annotate the masses of various motifs, such as anhydro- and deacetylation of the disaccharide backbone, non-proteogenic and amidated-amino acids to enable PGN search and analysis. In contrast, PGN_MS2 is specialized to predict MS/MS spectra for the PGN chemotypes, where the user simply selects the desired structural features of PGNs without needing to calculate and input their respective masses, rendering the analysis process user-friendly and flexible to accommodate novel modifications. Our algorithms encode the *in silico* fragmentation patterns of PGNs using rules derived from the experimental MS/MS spectra of known PGN standards. Unlike a simple fragmentation diagram in Byonic, PGN_MS2 generates the *in silico* predicted spectra of individual PGN precursor ions, thus facilitating spectral matching and scoring for PGN identification. To validate the reliability of our workflow, we compared the cosine similarity scores between the *in silico* predicted spectra of PGNs with the authentic spectra of several PGN motifs. Remarkably, PGN_MS2 consistently outperformed other spectral simulation software in metabolomics and proteomics, giving rise to higher similarity scores for PGN analysis. Moreover, the PGN_MS2 predicted spectra matched well with the fragmentation data acquired by different instruments (*i.e.,* Orbitrap and Q-TOF), showcasing the congruity of the *in silico* PGN fragmentation rules. Empowered by the PGN_MS2 library, we further demonstrated the facile and accurate assignment of closely related PGN isomers via automated spectral matching and scoring. However, we noted that the experimental MS/MS spectra of low abundant analytes tend to have lower quality, which led to the top predicted PGN structures having very close similarity scores. In these cases, manual inspections are needed to ensure accurate structural assignment. To facilitate such manual analysis, our PGN_MS2 records the precursor, fragmentation type, and chemical structure of all fragment peaks generated (**Figure S2D**).

During the preparation of our manuscript, Hsu *et al.* reported a high-throughput automated muropeptide analysis (HAMA) framework that generates *in silico* MS/MS fragments for PGN analysis.(96) However, we note several key distinctions between our workflow and HAMA. First, for *in silico* prediction of MS/MS patterns, HAMA focuses on fragmentation of the stem peptide; solely generating the b- and y- ions of stem peptides without any fragmentations of the sugar moieties in PGNs. HAMA restricts the types of PGN modifications to <6 (including those on sugar motifs and peptide aminations etc.) to avoid mass coincidences. On the other hand, our PGN_MS2 is developed especially for simulating MS/MS patterns of the soluble muropeptide chemotypes, whose fragmentation rules were derived from empirical analysis that include both for both sugar and peptide moieties in PGNs, showcasing superior matches to actual MS/MS data from HCD and CID fragmentations. As a result, our workflow accommodates much more diverse PGNs in the database and accurately distinguishes structural isomers by MS/MS matching. Notably, HAMA is reportedly unable to differentiate the 3-4 and 3-3 crosslinks in dimeric PGNs and hence can only consider 3-4 crosslinks currently. In contrast, with our PGN workflow, the 3-4 and 3-3 crosslinked PGN isomers can be facilely identified with signature fragments from the *in silico* MS/MS patterns (**Figure 4c and S7**). Moreover, our PGN_MS2 also includes specific fragmentations pertinent to the isoGln/Glu (q1/q2 and e1/e2), rendering its unique power in determining the amidation positions on isomeric PGNs (**Figure S3**). Regrettably, HAMA was shown to have difficulties in resolving these isomeric PGNs by MS/MS matching. Furthermore, PGN_MS2 creates a PGN MS library that is compatible with various vendors or open-source MS analysis software, offering users flexibility in choosing their preferred platforms for data analysis. To promote open-access research in the PGN field, PGN_MS2 itself is open source ([link](https://github.com/jerickwan/PGN_MS2)), where users can download directly to use or modify the code to increase the scope of fragmentations and the identities of amino acids and/or glycan motifs, *etc*. Recognizing the lack of PGNs in the existing metabolomics databank, we also uploaded the annotated MS/MS spectra of PGNs across different bacteria to the metabolomic data repository MoNA ([link](https://mona.fiehnlab.ucdavis.edu/spectra/browse?query=exists(tags.text:%27QiaoLab_PGN%27))). Expanding the collections of actual MS data for PGNs may greatly facilitate the future development of machine-learning-based *in silico* fragmentation modelling methods for PGN analysis.

Aided by PGN_MS2, we uncovered that the probiotic *Bifidobacterium spp.* showcases a large abundance of anNAM termini in peptidoglycan, which are non-hydrolytic cleavage products of LT enzymes.(81) By homology searching of known LTs, we identified and biochemically characterized three enzymes as the *bona fide* LTs in *Bifidobacterium*, namely, MltG, RpfB-FL, RpfB-truncated, respectively. Interestingly, MltG strictly requires nascent peptidoglycan strands as substrates for the non-hydrolytic cleavage, whereas RpfBs robustly use mature sacculi to produce anhydro-NAM termini. The complementary substrate preferences of these LTs may account for the remarkably high amount of anhydro-PGNs in *Bifidobacterium spp.* during normal growth. Given that *Bifidobacterium spp.* exhibit a characteristic Y-shaped rod morphology,(97) we speculate that the abundance of anNAM termini may be important for its cell shape maintenance since specific peptidoglycan features in bacteria are known to be important for cell morphology. For instance, the carboxypeptidase-trimmed stem peptides in peptidoglycan are critical for the helical shape of *Helicobacter pylori*.(98, 99) Future genetic studies to inactivate the putative LTs may offer more clues on the potential involvement of anhydro-NAM in *Bifidobacterium* cell shape determination. On the other hand, *Bifidobacterium spp.* are well-known probiotics that confer beneficial effects to hosts such as reducing LPS-induced inflammation *in vitro* and *in vivo*,(27, 28) we wondered if such predominant anhydro-PGN motifs may contribute to the anti-inflammatory effects of *Bifidobacterium spp.* Surprisingly, the pre-treatment with a synthetic anhydro-PGN effectively suppressed the LPS-induced proinflammatory cytokine expressions in murine macrophage RAW 264.7 cells *in vitro*. As *Bifidobacterium* anhydro-PGNs are non-agnostic to the canonical NOD1 and NOD2 immune receptors,(91–94) the underlying mechanisms of their anti-inflammatory roles are yet to be elucidated. Importantly, our work suggests the anti-inflammatory effects of *Bifidobacterium spp.* as probiotics likely are attributed to such abundant anhydro-PGNs. We are currently working to genetically manipulate the putative LTs in *Bifidobacterium spp.* to evaluate the anti-inflammatory activities of the mutants *in vivo*, which may lead to improved probiotics. Overall, our studies established novel and robust tools for comprehensive analysis of bacterial peptidoglycan structures, which will lead to fundamental biological discoveries in our emerging understanding of gut microbiota-derived PGNs in hosts.

## Acknowledgments

We acknowledge members of the Qiao lab for critical reading of the manuscript. J.M.C.K.’s PhD candidature is supported by the Nanyang Presidential Graduate Scholarship. This work was supported by the National Research Foundation (NRF) Singapore, NRF-NRFF12-2020-0006, NTU-Start-up grant, and MOE AcRF Tier 1, RG3/22 to Y.Q.

## Author Contributions

J.M.C.K. and Y.Q. designed the research; J.M.C.K. performed coding and developed the PGN analysis pipeline; J.M.C.K. and E.W.L.N performed biochemical characterization of Bifidobacterium LTs; Y.L. performed the *in vitro* assays with RAW264.7 and HEK-Blue reporter cells; Y.L., C.L, and Y.Z contributed to LC-MS method development; E.K. and S.W. provided anaerobic cultures of the gut bacteria; X.Z. and X.L. provided synthetic ah-PGN for testing; J.M.C.K., Y.L. and Y.Q analyzed data; S.W. and Y.Q. supervised the work; J.M.C.K. and Y.Q. wrote the paper with inputs from all authors.

